# A meta-analysis examining how fish biodiversity varies with marine protected area size and age

**DOI:** 10.1101/2023.03.27.534372

**Authors:** Helene A. L. Hollitzer, Felix May, Shane A. Blowes

## Abstract

Marine Protected Areas (MPAs) are a well-established conservation practice worldwide, but their effectiveness in protecting or replenishing fish biodiversity remains uneven. Understanding the patterns of this heterogeneity is central to general guidelines for MPA design and can ultimately provide guidance on how to maximize MPA potential. Here, we examine associations between the degree of protection, duration of protection, and protected area size, with fish biodiversity inside of protected areas relative to that of sites nearby, but outside of protected areas. We quantitatively synthesize 116 published estimates of species richness from 72 marine protected areas, and 38 estimates of Shannon entropy from 21 marine protected areas. We show that species richness is on average 18% (95% confidence intervals: 10% to 29%) higher in protected areas relative to areas open to fishing; and, on average Shannon entropy is 13% (95% confidence intervals: −2% to 31%) higher within protected areas relative to outside. We find no relationship between the degree or duration of protection with the ratio of species richness inside versus outside of protected areas; both fully and partially protected areas contribute to the accumulation of species inside protected areas, and protected areas of all ages contribute similarly on average to biodiversity conservation. In contrast to our expectations, increasing protected area size was associated with a decreased ratio of species richness sampled at sites inside versus outside the protected area, possibly due, for example, to insufficient enforcement and/or low compliance. Finally, we discuss why meta-analyses such as ours that summarize effect sizes of local scale biodiversity responses, that is, those at a single site, can only give a partial answer to the question of whether larger protected areas harbor more species than comparable unprotected areas.

## 1. Introduction

The world’s oceans are threatened by expanding anthropogenic impacts. Concerns are rising over observed declines in global fish stocks, as well as the extinction and redistribution of species (Arthington et al. 2016; Lam et al. 2020; Bijma et al. 2013). To counter these developments, Marine Protected Areas (MPAs) have become a major focus in ecosystem-based conservation and fisheries management (Gaines et al. 2010). However, while many studies indicate positive impacts of protection on biodiversity, effects remain uneven (Claudet et al. 2008; Watson et al. 2014). Some authors have suggested that the response of diversity to protection is a function of other factors, varying with the degree of protection, duration of protection, and the size of the protected area (Zupan et. al 2018; Claudet et al. 2008; Botsford, Micheli, and Hastings 2003; Ban et al. 2017).

The degree of protection varies among MPAs and is often expected to influence how communities respond to protection. No-take MPAs, also referred to as marine reserves or fully protected areas, have all extractive activities prohibited and are often associated with high conservation benefits (Sala and Giakoumi 2017; Grorud-Colvert et al. 2021). In the short term, however, fully protected areas often involve high socio-economic costs and are frequently confronted with opposition by extractive users (Lester and Halpern 2008). Partially protected areas vary in their regulations, allowing extractive activities to different degrees (e.g., prohibition of specific fishing practices such as commercial trawling or longline fishing), and are in some cases seen as politically more feasible, posing a balance between conservation and socio-economic viability (Lester and Halpern 2008; Sciberras et al. 2013; Zupan et al. 2018). A relatively recent meta-analysis by Zupan et al. (2018) found that the response to protection for finfish abundance to partial protection varied with different types of regulation. However, the impact of the degree of protection on biodiversity is less clear. Lester and Halpern (2008) found no significant relationship between the degree of protection (fully protected versus partial protected) and species richness, although this conclusion was limited by the low sample size of this study (n = 4).

MPAs have been hypothesized to show little or no change in biodiversity immediately following protection (Claudet et al. 2008; Halpern and Warner 2002; Soykan and Lewison 2015). This hypothesis is based on the premise that long-lived species, which generally exhibit long generation times, may take longer to show a response to protection. However, existing results are heterogeneous. Claudet et al. (2008) studied 80 fully protected areas (112 measurements) in a single temperate region (i.e., European reserves in the western Mediterranean Sea), and found support for the hypothesis, showing that protection effects on species richness increase with the duration of protection. In contrast, Halpern and Warner (2002) studied 19 fully protected areas (58 measurements) distributed across the globe and rejected the hypothesis as they found no relationship between protection effects on species richness and the time elapsed since establishment. Similarly, Soykan and Lewison (2015), who based their analysis on 50 data sets including fully and partially protected areas, found no effect of age of protected area and diversity inside versus outside of protected areas.

Protection may also affect biodiversity differently based on the size of the protected area. Modelling efforts aimed at fisheries management have suggested that larger MPAs should be more effective in restoring (or conserving) biodiversity by providing greater protection for highly mobile fishes and fishes undertaking migrations, and by allowing sufficient self-recruitment for species with long larval dispersal distances to be self-sustaining (Botsford, Micheli, and Hastings 2003). More fundamentally, larger protected areas often harbor more individuals (e.g., Claudet et al. 2008; Lester et al. 2009; Zupan et al. 2018), which is expected to result in more species via the more-individuals hypothesis (Srivastava and Lawton 1998). Encompassing more extensive areas, larger protected areas are also likely to contain a greater variety of habitats that offer suitable conditions for different species (Ban et al. 2017). While existing meta-analyses have shown that larger protected areas do often have higher densities of individuals (e.g., Claudet et al. 2008; Lester et al. 2009; Zupan et al. 2018), evidence for greater biodiversity benefits from larger reserves has been less prevalent. Neither a meta-analysis including partially and fully protected areas (Soykan and Lewison 2015) nor meta-analyses restricted to the results of fully protected areas (Halpern 2003; Claudet et al. 2008; Lester et al. 2009) found a significant correlation between MPA size and biodiversity indicators.

Here, we examine how the degree and duration of protection, and the size of the protected area impacts the effects of protection on fish biodiversity by collating studies from across the globe. By updating previous meta-analyses, we examine for generalities in the effectiveness of marine protected areas for biodiversity conservation, and identify sources of variation in outcomes. Understanding when and under which conditions MPAs are most effective forms an essential foundation for informing decisions about their establishment and governance. We examine both major types of protected areas, fully and partially protected areas in our analyses, offering decision-makers information that may have wider application. We aim to understand conservation effects on both the richness and evenness components of marine biodiversity, and thus avoid problems that emanate from considering a single diversity index (Roswell, Dushoff, and Winfree 2021). To meet our objective, we integrate results across studies, combining information from marine protected areas across the globe in an objective and quantitative way.

## 2. Methods

To synthesize results across studies we went through the steps of (1) source selection; (2) data acquisition and covariate extraction; (3) effect size calculation; and (4) statistical modelling. With this procedure we aim to reach generalizations about MPA effects on fish biodiversity, quantify their magnitude and variation, as well as to identify the sources of heterogeneity in results.

### 2.1 Source Selection

To compile a comprehensive database of studies that document the effects of protection on fish biodiversity, we searched the Web of Science on 15 April 2021 with following keywords (* represents a wildcard as any group of characters, including no character): (marine) AND (“marine protected area*” OR “marine reserve*” OR “no-take zone*” OR “fisher* closure*” OR “fully protected area*” OR “protected area” OR “marine sanctuary”) AND (biodiversity OR diversity OR “species richness”) AND (fish OR fishes).

Due to the wide range of terminology frequently used for MPAs, we designed the search string to be as comprehensive as possible. All candidate publications were subjected to a two-stage process in which we first filtered for suitable articles based on the article abstracts and, in a second stage, filtered the retained articles (e.g., if the abstract of an article provided insufficient information to exclude it) on the basis of the full texts. For a study to be included in our analysis, it must have satisfied the following criteria: (1) the study must have presented some quantification of biodiversity; (2) the study must have conducted a field assessment comparing a protected site to a reference site open to fishing (control-impact design), a site before and after MPA establishment (before-after design), or both (before-after, control-impact design); (3) the study must have measured fish biodiversity only, or reported biodiversity estimates for fish separately; and, (4) must have given information on the duration of protection, protected area size, and/or the name of the protected area (so as duration and size could be extracted from external databases).

In total, we got 1712 hits and screened the 649 candidate publications which we could access. We identified 38 unique studies that provided the required data for inclusion. We subjected all included studies to a critical appraisal of study quality. Following Sciberras et al. (2013) and Ohayon, Granot, and Belmaker (2021), we graded studies according to criteria further specified in Appendix S1: Table S2, and on this basis categorized them as being of high, medium, or low validity. About 29% of the studies resulted in high validity and about 71% in medium validity. No study was found to be of low validity (Appendix S1: Table S3).

### 2.2 Data Collection and Covariate Extraction

We extracted biodiversity estimates from inside and outside the MPA, before and after establishment, or both, from the text, tables, or figures of each publication. Numerical data from figures were extracted using WebPlotDigitizer (version 4.4). For each publication, we extracted data at the finest possible resolution for the following attributes: MPA (i.e., data from single MPAs were preferred over pooled data from multiple MPAs), depth, and habitat type (coral reef; rocky reef; sandy shores; open ocean; seagrass meadow; mangrove bay; kelp forest; macroalgae; hard bottom; soft bottom). For example, if a study reported data from multiple MPAs at different depths, all individual sets of data points were extracted and considered as unique samples and coded with the same study-specific ID assigning each sample to the respective study. Regarding the taxonomic level, we entered data in the most aggregated form because we were interested in the broadest response of fish diversity (e.g., if a study reported data on all fish, herbivores, and carnivores, we only extracted data for all fish). However, in some studies, data were only available for a subset of fish. This potentially affects the magnitude of effect, as different subsets of fish can be more or less species rich, and may respond differently to protection (e.g., fish targeted by commercial fisheries are assumed to benefit more from protection [Baskett and Barnett 2015]). To examine how this influenced our results, we categorized the data as documenting either fish targeted by commercial fisheries or fish not targeted by commercial fisheries. We chose this classification because it is commonly used and likely an important determinant of how fish respond to reduced or an end to exploitation. The categorization was preferably based on information from the respective paper. For studies where this information was not available, we used the Marine Trophic Index (mean trophic level of fisheries landings) of the study region (Pauly et al. 1998) to translate trophic level into susceptibility to exploitation. In studies where multiple time steps were reported, we extracted the most recent data as they represent the longest duration of protection; and, if several studies surveyed the same MPA, we retained biodiversity estimates of the most recent study for the same reason (unless the studies reported different biodiversity indices). We found only seven studies with data from before the onset of protection (before-after: n = 2; before-after, control-impact: n = 5; Appendix S1: Table S1), preventing an analysis of before-after data. We therefore discarded all before data and exclusively retained the most recent data for analyses. As a result, before-after, control-impact studies were equated to control-impact studies, and before-after studies were omitted from further analyses because no suitable control was available.

Second, we extracted data on three covariates characterizing each MPA: degree of protection (fully vs. partially protected), duration of protection (years between MPA establishment and date of survey) and protected area size (area in km^2^). We extracted data on these characteristics for each MPA from the text, tables, and figures of each publication. One study combined fully and partially protected areas into a single effect size (Friedlander, Brown, and Monaco 2007), which we treated as partially protected, as the estimate derived from varying degrees of restrictions and thus better fit the ‘partially protected’ category. When biodiversity estimates of multiple MPAs were exclusively provided in an aggregated form, the mean value of duration of protection and protected area size was used as the corresponding age and size value. If the size was not specified in the text or a table but a map, size was extracted via the image processing program ImageJ (version 1.53k). ImageJ enables calculation of areas by setting the scale using the map scale bar and manually marking out the MPA area. In cases where specifications on age or size were not provided or otherwise available from the study, the information was obtained from the World Database on Protected Areas (UNEP-WCMC and IUCN 2021).

### 2.3 Effect Size Calculation

From the data, we calculated the biodiversity effect sizes as the log response ratio, calculated as the natural logarithm of the ratio of mean diversity at the protected site 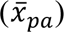 over the mean diversity at the unprotected (open to fishing) site 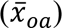 (Hedges, Gurevitch, and Curtis 1999):

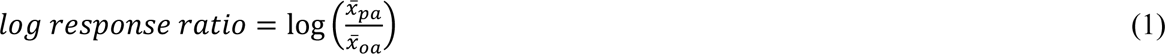

The log response ratio has an approximate normal distribution (Hedges, Gurevitch, and Curtis 1999), and positive effect size values indicate greater diversity inside the MPA boundaries relative to outside, whereas negative values indicate greater diversity outside the MPA boundaries relative to inside.

### 2.4 Statistical Models

First, to quantify the overall effect of protection on fish biodiversity, we fit separate models for the log response ratio of species richness and of Shannon entropy (Table 1: Model 1 and 7); all other biodiversity metrics had sample sizes too small for statistical analysis (Appendix S2: Table S1). All models fit allowed effect sizes to vary among studies by including a study-specific ID as a random intercept, and we first estimated the overall average across all studies. Second, we examined whether the log response ratio of diversity varied with three MPA characteristics. To do this, we fit mixed-effects models to quantify the relationship between the log response ratio and the degree of protection (fully vs. partially protected area), the duration of protection and protected area size; both duration and size were log-transformed and mean-centered before model fitting. We also examined whether the response to protection varied between fish subgroups with different susceptibility to exploitation by including a categorical covariate with the levels fish targeted by commercial fisheries and fish not targeted by commercial fisheries; study-specific ID was included as a random intercept in all models. We also examined two-way interactions between all covariates. We fit models using restricted maximum likelihood and compared models using the corrected Akaike Information Criterion (AICc; Burnham and Anderson 2002). Visual inspection of residual diagnostic plots did not reveal obvious deviations from homoscedasticity or normality. In case of doubt, we removed critical data points to verify whether the results remained constant, which was true in all cases. The significance of fixed effect model parameters was assessed using Satterthwaite’s method for calculating degrees of freedom as provided in the R package lmerTest (version 3.1-3, Kuznetsova, Brockhoff, and Christensen 2017).

**Table 1.** Model specifications and corresponding AICc values.

| No. | Model | df | Obs. | Gr. | AICc |
| --- | --- | --- | --- | --- | --- |
| 1 | $\ln\text{RR}(\text{SR}) \sim 1 + (1 \mid \text{study ID})$ | 3 | 116 | 35 | 68.16169 |
| 2 | $\ln\text{RR}(\text{SR}) \sim \text{FG} + \text{size} + (1 \mid \text{study ID})$ | 6 | 116 | 35 | 69.00059 |
| 3 | $\ln\text{RR}(\text{SR}) \sim \text{FG} + \text{age} + (1 \mid \text{study ID})$ | 6 | 116 | 35 | 77.25142 |
| 4 | $\ln\text{RR}(\text{SR}) \sim \text{FG} + \text{protection} + (1 \mid \text{study ID})$ | 6 | 116 | 35 | 74.96522 |
| 5 | $\ln\text{RR}(\text{SR}) \sim \text{FG} + \text{protection} + \text{age} + \text{size} + (1 \mid \text{study ID})$ | 8 | 116 | 35 | 81.52182 |
| 6 | $\ln\text{RR}(\text{SR}) \sim (\text{FG} + \text{protection} + \text{age} + \text{size})^2 + (1 \mid \text{study ID})$ | 17 | 116 | 35 | 114.15522 |
| 7 | $\ln\text{RR}(\text{H}) \sim 1 + (1 \mid \text{study ID})$ | 3 | 38 | 11 | 32.83216 |

**Abbreviations:** df = number of parameters in the model; Obs. = number of observations; Gr. = number of studies; $\ln\text{RR}$ = log response ratio; SR = species richness; study ID = unique identifier for each study; FG = fish group as all, commercially targeted, and non-commercial fish; size = mean-centered and log-transformed MPA size in $\text{km}^2$ ; age = mean-centered and log-transformed MPA age in years; protection = protection degree as fully protected and partially protected; H = Shannon entropy

In addition, we fit meta-analytic models to data where we had estimates of the effect size uncertainty. Since the results of the two approaches (without and with estimates of effect size uncertainty) were not substantially different (Appendix S2: Table S2 and Figs. S4-S5), we report results using the analysis without effect size uncertainty. This allowed more studies to be retained and potentially produces more precise results than an analysis using a restricted database of estimates with standard errors (Morrissey 2016).

To aid interpretation, mean effect sizes and confidence intervals (CIs) are presented in the text as percent changes calculated as % *change* = 100 · [exp (*lnRR*) − 1] (Pustejovsky 2018). Regression slopes were translated into the change in the response associated with a ten-fold increase in the covariate of interest, and similarly converted into a percent change. We conducted all analyses in the R (version 3.6.3, R Core Team 2020) using the R packages lme4 (version 1.1-26, Bates et al. 2015), lmerTest (version 3.1-3, Kuznetsova, Brockhoff, and Christensen 2017), MuMIn (version 1.43.17, Bartoń 2020), and ggplot2 (version 3.3.3, Wickham 2016).

## 3. Results

### 3.1 General Description of the Compiled Database

The database of species richness results contained 35 independent studies, 77% of which presented species richness as the only measure of biodiversity (Appendix S2: Table S1). The papers were published between 1996 and 2021; 71% of which were published in the last decade between 2011 and 2021. The database documented 59 MPAs individually sampled, and 13 groupings ranging from 2 to 13 protected areas (i.e., the results of several MPAs were pooled in the original study). In total, we extracted 116 different sets of biodiversity estimates from the studies, as some studies examined, for example, multiple fish groups such as commercial vs. non-commercial or multiple habitat types within one MPA.

The data originated from globally distributed MPAs spanning the Pacific, Atlantic, and Indian Ocean, the Mediterranean Sea, and included both temperate and tropical regions (Fig. 1a). Of the 116 effect size calculations (i.e., log response ratios), 71 compared biodiversity of an unprotected area with that of a fully protected area, and 45 compared an unprotected area with a partially protected area. MPAs were protected for an average of 14.4 years (median = 10 years), with the oldest being 52 years old and the youngest 1.5 years old (Fig. 1b). The average size of the 72 MPAs or MPA groupings was 167.4 km^2^ (median = 9.7 km^2^), with the largest MPA being 3337 km^2^ and the smallest 0.008 km^2^ (Fig. 1c).

**Figure 1.**
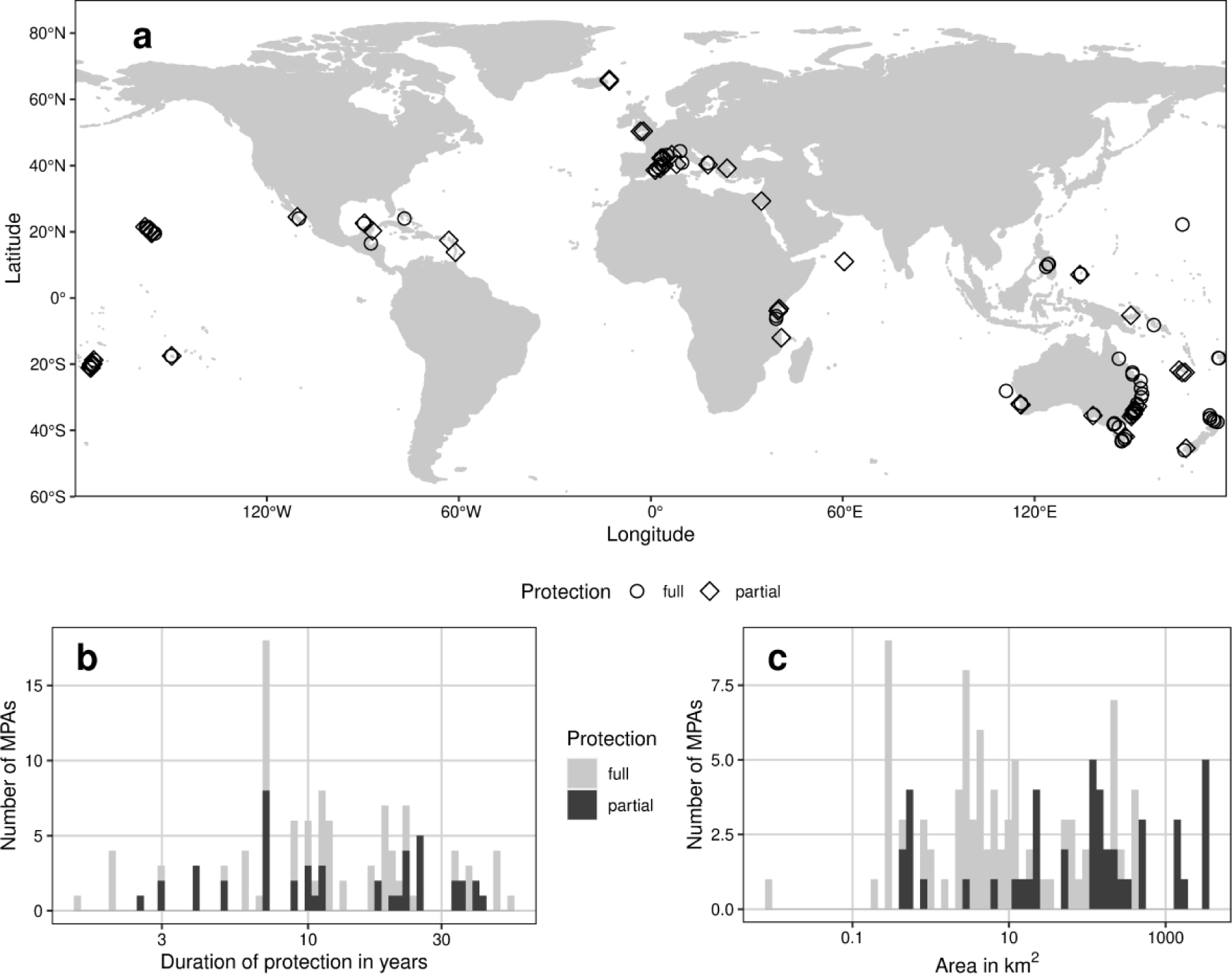
(a) Map of included MPAs. Since in some studies the biodiversity estimates of several MPAs were aggregated into a single value, not every point corresponds to a separate log response ratio entry. (b) and (c) show the distribution of the analyzed MPA characteristics, subdivided into (b) MPA age as the number of years between MPA establishment and date of survey and (c) MPA size in km^2^ and. MPA age and size both are binned on a log scale. The histograms are based on the database of species richness results.

From the 11 papers (published between 2002 and 2021) presenting Shannon entropy we extracted 38 sets of biodiversity estimates.

### 3.2 Quantitative Analysis

#### 3.2.1 Effect of protection

Overall, species richness was significantly higher inside MPAs relative to sites open to fishing. Species richness was on average 18.4% (95% CIs: 10.2% to 27.5%) higher within MPA boundaries when compared to adjacent unprotected sites (p = 0.0002; Fig. 2a; Table 2: Model 1). Shannon entropy was on average 13.4% (95% CIs: −1.59% to 30.5%) higher in MPAs than in adjacent unprotected sites (p > 0.05; Fig. 2b; Table 2: Model 7).

**Figure 2.**
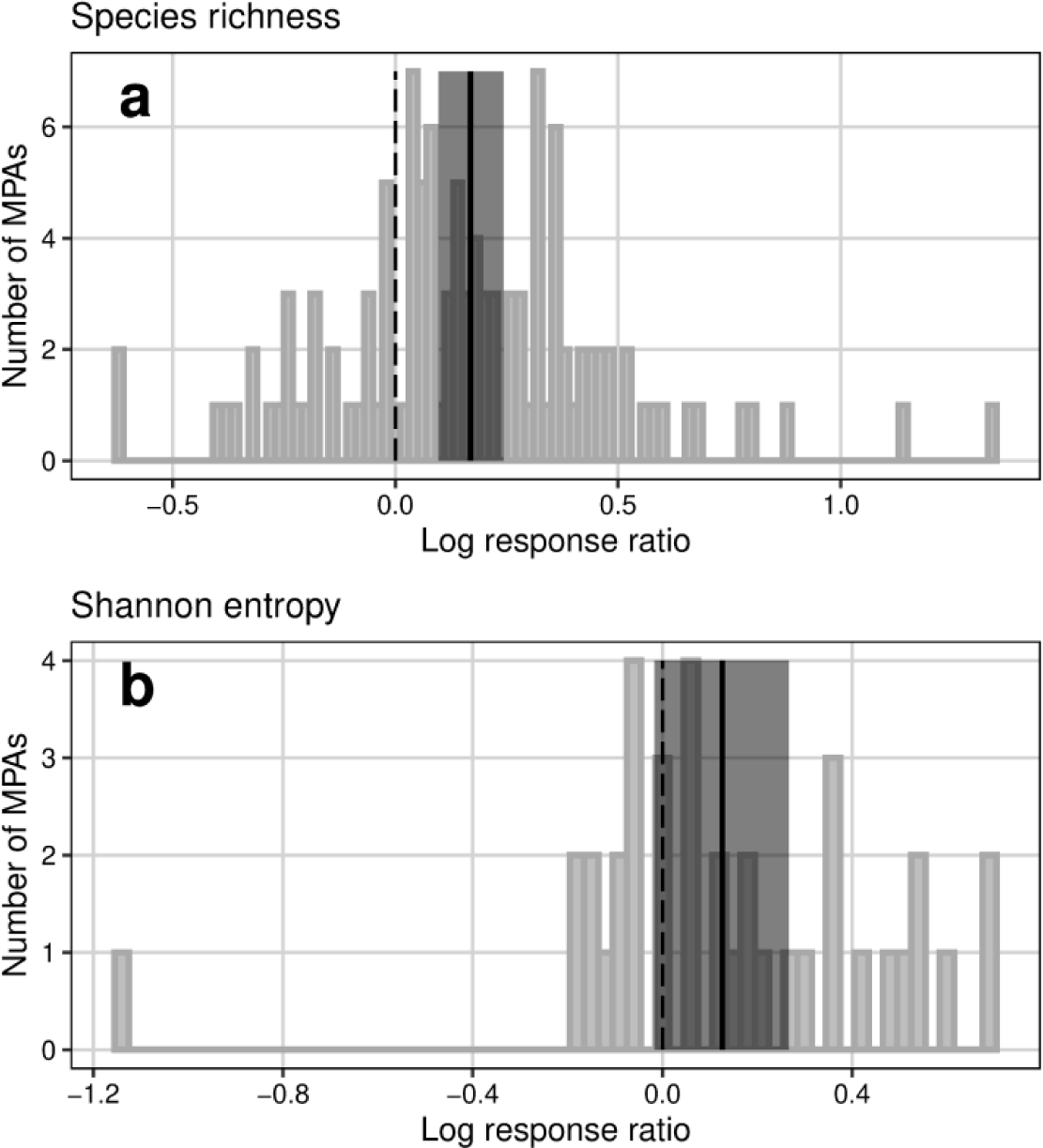
Overall effect of protection on biodiversity shown as the distribution of log response ratios (MPA:unprotected) of (a) species richness and (b) Shannon entropy. The average MPA efficacy in protecting biodiversity as the mean log response ratio of species richness and Shannon entropy are depicted as solid black lines and 95% confidence intervals are indicated with grey shading. The corresponding values can be found in Table 2: Model 1 and 7 for (a) and (b), respectively. In both subfigures, a log response ratio of 0 indicates no effect through protection. Positive log response ratios indicate greater diversity inside the MPA boundaries relative to outside and negative values a greater diversity outside the MPA boundaries relative to inside. The sample sizes for (a) and (b) were 116 and 38, respectively.

**Table 2.** Summary of models fits to log response ratios.

| Model<sup>3</sup> | Effect | Estimate | 95% Confidence Interval |  | p-value |
| --- | --- | --- | --- | --- | --- |
|  |  |  | *Lower* | *Upper* |  |
| 1 | Intercept | 0.169 | 0.097 | 0.243 | 0.00021 |
| 2 | Intercept | 0.120 | 0.028 | 0.211 | 0.02042 |
|  | commercial fish | 0.286 | 0.065 | 0.508 | 0.01866 |
|  | non-commercial fish | 0.003 | -0.220 | 0.227 | 0.97838 |
|  | MPA size | -0.046 | -0.073 | -0.017 | 0.00125 |
| 3 | Intercept | 0.151 | 0.067 | 0.234 | 0.00275 |
|  | commercial fish | 0.225 | 0.004 | 0.442 | 0.04695 |
|  | non-commercial fish | -0.065 | -0.283 | 0.152 | 0.55578 |
|  | MPA age | -0.018 | -0.104 | 0.063 | 0.66202 |
| 4 | Intercept | 0.187 | 0.083 | 0.294 | 0.00200 |
|  | commercial fish | 0.239 | 0.016 | 0.466 | 0.04610 |
|  | non-commercial fish | -0.057 | -0.281 | 0.172 | 0.62810 |
|  | Partial protection | -0.086 | -0.213 | 0.052 | 0.18350 |
| 7 | Intercept | 0.126 | -0.016 | 0.266 | 0.103 |

**Note:** Models 5 and 6 are not presented as only results from the model with the lowest AICc that included the variable of interest were reported.
<sup>3</sup> Model numbers refer to the models described in Table 1.

#### 3.2.2 Correlates of response to protection

We tested potential predictor variables (i.e., size; age; degree of protection) to elucidate the variability in effect sizes. All pairwise-interactions increased AICc values (Table 1), indicating no support for interacting effects of size, age, and degree of protection on the log response ratio. Model selection supported the inclusion of the fish group variable, which adjusts for studies that only examine a subgroup of fish that may benefit from protection to a lesser or greater degree than the mean (Table 2), with commercial fish benefiting protection more strongly in all models (Table 2; Appendix S2: Fig. S1). Model selection also showed that the model that included protected area size had a similar predictive power as the intercept only model (*Δ*AICc < 2), whereas all other models were not well supported by the data and showed increases in AICc of > 6 (Table 1).

The linear mixed model revealed a negative relationship between the effect size and size of MPAs for species richness, showing a slight but significant reduction relative to unprotected sites as MPA size increased (slope = −0.046; 95% CIs: −0.073 to −0.017; p = 0.001; Fig. 3a; Table 2: Model 2). The slope suggests an average decrease in species richness relative to unprotected areas of 10.1% for every ten-fold increase in MPA size. The log response ratio of species richness was not significantly related to the duration of protection (slope = −0.018; 95% CIs: - 0.104 to 0.063; p > 0.05; Fig. 3b; Table 2: Model 3). Similarly, there was no difference between fully protected and partially protected areas on the species richness log response ratio (slope = - 0.086; 95% CIs: −0.213 to 0.052; p > 0.05; Table 2: Model 4).

**Figure 3.**
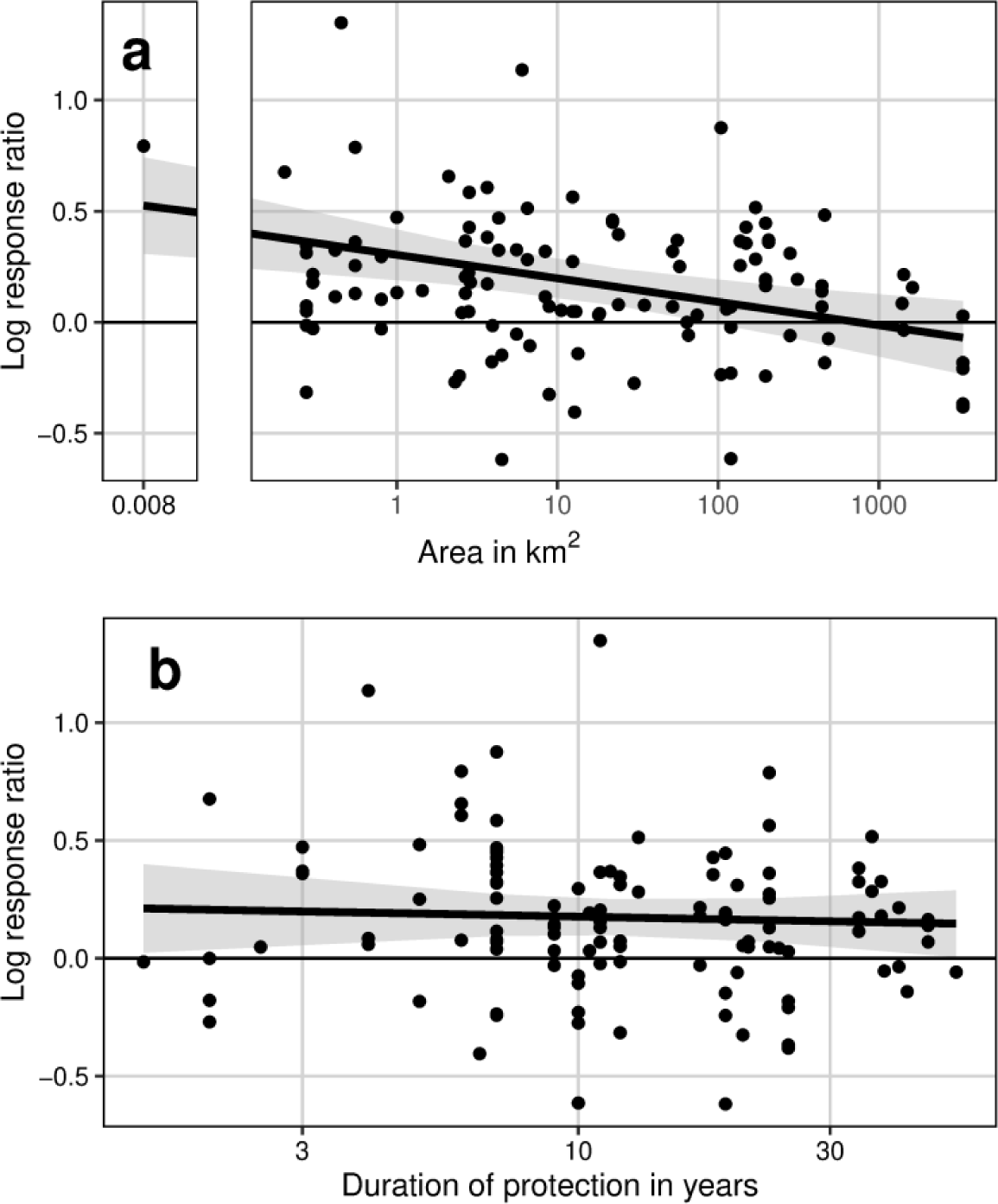
Log response ratio (MPA:unprotected) of species richness as a function of (a) MPA size in km^2^ and (b) MPA age as the number of years between MPA establishment and date of survey. Both x-axes are on a log-scale and 95% confidence intervals are indicated with grey shading. Corresponding values can be found in Table 2: Model 2 and 3 for (a) and (b), respectively. In both subfigures, a log response ratio of 0 indicates no effect through protection. Positive log response ratios indicate greater diversity inside the MPA boundaries relative to outside and negative values indicate greater diversity outside the MPA boundaries relative to inside. The sample size for (a) and (b) was 116 each. Note: in (a), the plot is split into two subplots for better readability of the data.

## 4. Discussion

Using 116 comparisons compiled from 35 studies, we find that protected areas are on average associated with an 18.4% increase in species richness when compared to adjacent unprotected sites. Contrary to our expectation that larger MPAs would have comparatively more species relative to adjacent unprotected areas, we found a negative relationship between MPA size and the log response ratio of species richness. We found no relationship between the time elapsed since implementation or the degree of protection with the effectiveness of MPAs. We also found a weaker response in the log response ratio of Shannon entropy, which was 13.4% higher in MPAs than in unprotected areas, an increase that was not statistically significant, albeit with a smaller sample size (n = 38 comparisons).

Contrary to the prediction that larger protected areas are more effective in protecting and restoring biodiversity, we found smaller effect sizes for species richness as protected area size increased. There are several possible factors that could be contributing to this result. Enforcement of regulations and compliance, both of which are critical factors in ensuring that protected areas achieve their designated objectives (Walmsley and White 2003; De Santo 2013; Fujitani et al. 2012; Rife et al. 2012) are likely harder in large marine protected areas (Rife et al. 2012; De Santo 2013; Pala 2013; Wilhelm et al. 2014). For example, large-scale protected areas have been shown to be less respected by locals with illegal fishing and poaching sometimes remaining after protection (De Santo 2013). Such compliance related issues emerge especially in intensively used areas (Wilhelm et al. 2014; Roberts et al. 2003), and when no alternative sources of income are provided (Mora and Sale 2011). To minimize conflicts with stakeholders, large marine protected areas are therefore often established from the outset in remote regions with less commercial and recreational fishing pressure, while smaller protected areas tend to be sited in coastal areas with high commercial interest (Devillers et al. 2014; Devillers et al. 2020), likewise restricting potential conservation value (Mizrahi et al. 2018).

The number of large marine “paper parks” (i.e., MPAs that exist on paper but are not properly regulated) has increased in recent years due to rising global pressure to protect large percentages of the ocean. However, many of these MPAs are implemented without sufficient resources (Rife et al. 2012; De Santo 2013; Jones and De Santo 2016). The focus on protecting a certain percentage of the ocean might encourage the prioritization of political over ecologically driven decision-making (Agardy, Di Sciara, and Christie 2011; De Santo 2013), leading to MPAs lacking adequate quality to achieve designated conservation goals (Barnes et al. 2018).

Size effects of MPAs have also been hypothesized to only occur for very large MPAs (i.e., > 500 km^2^; Halpern 2003). Here, the median size of protected areas was just under 10 km^2^. While this is a larger median size compared to prior meta-analyses (e.g., Halpern 2003 [median size = 4.0 km^2^]; Claudet et al. 2008 [median size = 1.95 km^2^]; Lester et al. 2009 [median size = 3.3 km^2^]), sizes included in our analysis might still be insufficient to detect a strong influence of size if the threshold for an effect to occur was even higher. And although our database reflects the global database on protected areas reasonably well, very large MPAs are not represented in the sample, especially for fully protected areas (Appendix S2: Fig. S2).

Against the expectation that older protected areas would yield greater biodiversity conservation, we found that MPAs were equally effective across all ages. This is a similar finding to Halpern and Warner (2002), who found that effects driven by protection manifest within the first three years after implementation, thus making an effect unable to be detected by the linear models applied in this study. Our analysis contained 9 MPAs protected for ≤ 3 years. Looking at the respective data points, of the 6 MPAs that have been under protection for 1 to 2.5 years, 5 fell below the log response ratio predicted for their respective age. The 3 MPAs that have been protected for 3 years all fell above the log response ratio estimated for that age. The fitting of further non-linear models was omitted, as the small sample size would have precluded reliable results. Molloy, McLean, and Côté (2009) also found that large species did not respond slower to protection in terms of their density. They hypothesized that the larger home ranges of large and long-lived species meant that they were more likely to encounter and settle in newly established MPAs. Similar mechanisms may also play a role in the impact of protection on biodiversity, masking the importance of life-history traits.

Whether partially protected areas are helping global conservation is subject of much debate. We found, congruent with Giakoumi et al. (2017) and Lester et al. (2009), a non-significant trend towards fully protected areas being more effective than partially protected areas in protecting species richness. This corroborates the claim of several researchers, that, depending on the specific aims, both types of protected areas can make important contributions to nature conservation (Singleton and Roberts 2014; Jones and De Santo 2016; Devillers et al. 2020). We note, however, that our classification system may have been too coarse to establish a clear relationship between degree of protection and effectiveness. In particular, partially protected areas vary greatly in regulations, which is likely a major contributor to the heterogeneity of our results. Further work to disentangle how different regulations impact biodiversity conservation may need a more sophisticated classification system, such as those proposed by Grorud-Colvert et al. (2021) and Zupan et al. (2018).

Meta-analyses such as ours can only give partial answers to questions associated with diversity, such as whether larger or older protected areas harbor more species than comparable unprotected areas. This is because diversity, and its response to drivers is inherently scale-dependent (Chase et al. 2018). Whilst we did not observe any relationship between the grain of samples and effect sizes in the data we compiled (Appendix S2: Fig. S3), comparing local sites inside and outside of protected areas (i.e., the response of diversity at the local scale) is only part of the picture for assessing how diversity responds to protection in large areas. Analyses of local scale responses such as ours are best suited to capture predicted benefits for sites in larger protected areas due to improved demographic processes, such as increased survival, greater recruitment, and higher fecundity (Botsford, Micheli, and Hastings 2003). In addition, however, the null expectation is that with increasing area, environmental and habitat heterogeneity increases also, leading to greater species richness (Rosenzweig 1995; Ban et al. 2017). This means that assessments of the combined diversity of multiple sites inside large protected areas, encompassing the site-to-site variation of species composition (i.e., beta-diversity), are needed to fully evaluate their conservation value. Such analyses are often beyond the scope of meta-analyses, which are limited to published results. Even where multiple effect sizes from different sites within a given protected area are available, the (raw) data needed to combine (effort-standardized) diversity across sites are typically unavailable. To examine the scale-dependence of how diversity responds to protection, future studies should make use of increasingly available raw data that document species abundances across sites inside and outside of protected areas (see e.g., Blowes et al. 2020). This also permits the assessment of multiple biodiversity measures, including compositional changes (Hillebrand et al. 2017), and for teasing apart the contribution of how the total and relative abundance of individuals and species, and within species aggregation combine to determine the response of diversity to protection (Chase et al. 2018, Blowes et al. 2020).

## 5. Conclusions

This study demonstrates that protected areas across all ages and protection regimes can make a valuable contribution to biodiversity conservation. We argue that the finding that larger marine protected areas do not achieve the expected higher conservation impact should impel further investigation into the underlying causes (biological, political, and methodological), rather than the omission of large protected areas from future MPA planning. For that, a more holistic picture of biodiversity is needed. More studies are needed that (a) quantify beta-diversity and (b) use a range of indices rather than just species richness in order to embrace taxonomic, phylogenetic, and functional diversity. This will help to understand biodiversity responses to protection in more detail, prevent misinterpretations and enable improvements in MPA design. We propose that more attention should be devoted to suitable and (cost-)effective monitoring methods (Rowlands et al. 2019), and the incorporation of human wellbeing into marine protected area planning, as locals are directly and often heavily affected by their implementation (Rife et al. 2012; Fujitani et al. 2012; De Santo 2013; Rees et al. 2013). Only then can marine protected areas become sustainable both socially and ecologically (Kamat 2014).

## Acknowledgements

SAB acknowledges the the support of the German Centre of Integrative Biodiversity Research (iDiv) Halle-Jena-Leipzig (funded by the German Research Foundation; FZT 118).

## Author Contributions

Helene A. L. Hollitzer, Shane A. Blowes and Felix May conceived the ideas and designed the methodology. Helene A. L. Hollitzer collected and analyzed the data and led the writing of the manuscript, with additional contributions form Shane A. Blowes. All authors contributed to the interpretation of the results and to the editing of the manuscript into its final draft.

## Conflict of interest statement

The authors declare no conflict of interest.

## Notes

https://doi.org/10.5281/zenodo.7772379

### Competing Interest Statement

The authors have declared no competing interest.

