## Appendix S1 for "A meta-analysis examining how fish biodiversity varies with marine protected area size and age"

**Table S1** Hierarchy of study quality based on sampling design (Sciberras et al. 2013).

| **Sampling Design** | **Spatial Replication** | | **Temporal Replication** | | **Number of Studies** | **Total** | **Percent** |
|---|---|---|---|---|---|---|---|
| | *Treatment (MPA)* | *Control* | *Before* | *After* | | | |
| BACI | Multiple | Multiple | Multiple | Multiple | 2 | 5 | 12.50% |
| | Multiple | Multiple | Once | Multiple | 1 | | |
| | One | One | Once | Multiple | 2 | | |
| CI | Multiple | Multiple | - | Multiple | 4 | 33 | 82.50% |
| | Multiple | Multiple | - | Once | 14 | | |
| | One | Multiple | - | Multiple | 2 | | |
| | Multiple | One | - | Once | 1 | | |
| | One | Multiple | - | Once | 4 | | |
| | One | One | - | Multiple | 2 | | |
| | One | One | - | Once | 5 | | |
| BA | One | - | Multiple | Multiple | 1 | 2 | 5.00% |
| | One | - | Multiple | Once | 1 | | |

**Note:** Quality regarding sampling design decreases down the list.

**Table S2** Quality assessment based on Ohayon et al. (2021): appraisal criteria.

| Factor | High validity | Medium validity | Low validity |
|---|---|---|---|
| **Study design** | Equal number of sites inside and outside the MPA | Sites inside and outside the MPA; but not equally | Sampling design includes sites only inside or outside the MPA |
| | Temporal replication | No temporal replication | No temporal replication |
| | Spatial replication (MPA and control site) | Spatial replication (MPA or control) | No spatial replication |
| **Sampling** | Large sample sizes at each site; n >= 6 | Medium sample sizes at each site; n >= 3 | Small sample sizes at each site; n < 3 |
| | Habitat type at unprotected area and MPA are similar | Habitat type at unprotected area and MPA are medium similar; habitats not described in sufficient detail | Habitat type at unprotected area and MPA are unsimilar or not reported |
| **Accounting for potential effect modifiers** | MPA enforcement is high | MPA enforcement is sufficient; limited illegal fishing | MPA enforcement is low or not reported |

**Table S3** Quality assessment based on Ohayon et al. (2021): quality assessment of included studies.

| Study | Study design | Sampling | Accounting for potential effect modifiers | Estimated overall validity |
|---|---|---|---|---|
| **Mitchell et al. 2021** | no information on number of sites; spatial replication (MPA only); no temporal replication | unsimilar habitats; large sample sizes | consultation with local stakeholders identified the protected area as a currently managed area with greater protection | medium |
| **Turnbull, Johnston, and Clark 2021** | equal number of sites; spatial replication (MPA and control); no temporal replication | similar habitats; large sample sizes | enforcement is present, but low | high |
| **Favoretto et al. 2020** | unequal sites; no spatial replication; no temporal replication | similar habitats; large sample sizes | weak regulations | medium |
| **Smallhorn-West et al. 2020** | unequal sites; spatial replication (MPA and control); no temporal replication | similar habitats; large sample sizes | enforcement not reported | medium |
| **Ramirez-Ortiz et al. 2020** | unequal sites, spatial replication (MPA and control); temporal replication | habitat similarity not reported; large sample sizes | enforcement not reported | medium |
| **Ortodossi et al. 2019** | unequal sites; spatial replication (MPA and control); no temporal replication | similar habitats; large sample sizes | well-enforced | high |
| **Gress et al. 2018** | unequal sites; no spatial replication; no temporal replication | similar habitats (same coral communities, distinct health conditions); large sample sizes | enforcement not reported | medium |
| **Alonso Aller, Jiddawi, and Eklöf 2017** | equal sites; spatial replication (MPA and control); temporal replication | similar habitats; large sample sizes | one MPA well-enforced, one less strong | high |
| **Edgar et al. 2017** | unequal sites; spatial replication (MPA and control); no temporal replication | similar habitats; large sample sizes for most (10 of 12) | well-enforced | high |
| **McClanahan and Muthiga 2017** | unequal sites; spatial replication (control only); temporal replication | similar habitats (similar hard coral cover, similar number of coral genera; similar macroalgae | enforcement restricted to a few reefs | medium |

| | | | | |
|---|---|---|---|---|
| | | cover); large sample sizes | | |
| **Powell et al. 2016** | equal number of sites; spatial replication (MPA and control); temporal replication | habitat similarity not reported; large sample sizes | active enforcement | high |
| **Bertucci et al. 2016** | equal number of sites; spatial replication (MPA and control); no temporal replication | similar habitats; medium sample sizes | enforcement not reported | medium |
| **Smith and Anderson 2016** | equal sites for 2 of 3 MPAs - control pairs (6/6, 6/6, 5/6); spatial replication (MPA and control); temporal replication | similar habitats; large sample sizes | enforcement not reported | medium |
| **Humphries, McQuaid, and McClanahan 2015** | equal sites; spatial replication (MPA and control); no temporal replication | medium similar habitats; large sample sizes | enforcement not reported | medium |
| **Hehre and Meeuwig 2015** | unequal sites; spatial replication (MPA and control); no temporal replication | medium similar habitats; large sample sizes | well-enforced | high |
| **Alemu and Jahson 2014** | unequal sites; spatial replication (control only); no temporal replication | medium similar habitats; large sample sizes | enforcement not reported | medium |
| **Guidetti et al. 2014** | unequal sites; spatial replication (MPA and control); no temporal replication | similar habitats | well-enforced no-take MPAs, other categorized as PPAs | high |
| **Williamson et al. 2014** | unequal sites; spatial replication (MPA and control); temporal replication | medium similar habitats; large sample sizes | enforcement not reported | medium |
| **Sheehan et al. 2013** | unequal sites; no spartial replication; temporal replication | similar habitats; large sample sizes | enforcement not reported | medium |
| **Wind and Jack 2013** | unequal sites; spatial replication (MPA and control); temporal replication | habitats unsimilar; large sample sizes | enforcement not reported | medium |
| **Olds et al. 2013** | equal sites for 2 of 3 MPAs sampled; spatial replication (MPA and control); no temporal replication | similar habitats; large sample sizes | enforcement not reported | medium |
| **Rasher, Hoey, and Hay 2013** | equal sites; spatial replication (MPA and | unsimilar habitats; large sample sizes | well-enforced and high compliance | high |

| | | | | |
|---|---|---|---|---|
| | control); no temporal replication | | | |
| **Carassou et al. 2013** | unequal sites; spatial replication (MPA and control); no temporal replication | unsimilar habitats; large sample sizes | well-enforced | medium |
| **Noble et al. 2013** | equal sites; no spatial replication; no temporal replication | unsimilar habitats; large sample sizes | paper directs reader to 3 papers that sampled the same locations for site description in which is stated that enforcement and compliance are high/sufficient | medium |
| **Tessier et al. 2013** | equal number of sites; spatial replication (control only); no temporal replication | similar habitats; large sample sizes | fishing is "regulated" | medium |
| **Olds et al. 2012** | unequal sites; spatial replication (control only); no temporal replication | similar habitats; medium sample sizes | enforcement not reported | medium |
| **Coll et al. 2012** | equal sites; spatial replication (MPA and control); temporal replication | similar habitats; large sample sizes | enforced, no details | high |
| **Huntington et al. 2010** | unequal sites; no spatial replication; no temporal replication | medium similar habitats; large sample sizes | enforced, no details | medium |
| **Page et al. 2009** | equal number of sites; spatial replication (MPA and control); no temporal replication | similar habitats; medium sample sizes | 2 of 3 MPAs are well managed | medium |
| **Seytre and Francour 2009** | unequal sites; spatial replication (control only); temporal replication | similar habitats; large sample sizes | enforcement not reported | medium |
| **Edgar and Stuart-Smith 2009** | number of sites not given in sufficient detail; spatial replication (MPA and control); no temporal replication | habitat similarity not reported; large sample sizes | enforcement not reported | medium |
| **Harborne et al. 2008** | equal sites for one habitat and unequal sites for a second habitat; no spatial replication; no temporal replication | similar habitats; large sample sizes | well-enforced | medium |
| **Friedlander, Brown, and Monaco 2007** | no information on number of sites; spatial replication (MPA and control); no temporal replication | similar habitats; large sample sizes | enforcement not reported | medium |

| | | | | |
|---|---|---|---|---|
| **Claudet et al. 2006** | unequal sites: no spatial replication; temporal replication | similar habitats; large sample sizes | active enforcement | high |
| **Jaworski, Solmundsson, and Ragnarsson 2006** | equal sites; spatial replication (MPA and control); temporal replication | medium to unsimilar habitats; medium sample sizes | state that enforcement is present | medium |
| **Jones et al. 2004** | equal sites; no spatial replication; temporal replication | similar habitats; medium sample sizes | enforcement not reported | medium |
| **Khalaf and Kochzius 2002** | unequal sites; spatial replication (control only); no temporal replication | similar habitats; large sample sizes | occasionally illegal fishing appears | medium |
| **Russ and Alcala 1996** | equal sites; no spatial replication; temporal replication | similar habitats; large sample sizes | marine management committee overseeing successful protection | high |
