## Appendix S2 for "A meta-analysis examining how fish biodiversity varies with marine protected area size and age"

**Table S1** Information on the biodiversity indices used in the studies that met the inclusion criteria.

| **Index** | **Number**[1] **(only)**[2] | **Description** | **Formula** |
|---|---|---|---|
| Species Richness | 35 (27) | Count of different species represented in an area. | $SR = \sum_{i=1}^{S} p_i^0$ |
| Shannon Entropy | 11 (2) | Entropy that quantifies the uncertainty associated with predicting the outcome of a sampling process. The more species present in an area and the closer their relative abundances, the more difficult it is to correctly predict the outcome. | $H = -\sum_{i=1}^{S} p_i \ln p_i$ |
| Pielou's Evenness | 2 (0) | Measure of the equal distribution of individuals in a sample among species. The value lies between 0 (absolute unequal distribution) and 1 (uniform distribution). | $J = (-\sum_{i=1}^{S} p_i \ln p_i) / \ln(S)$ |
| Simpson's Diversity Index | 1 (0) | Probability that two individuals sampled randomly from a community are of the same species. | $\lambda = \sum_{i=1}^{S} p_i^2$ |
| Taxa Richness | 1 (0) | Count of different taxa represented in an area. | $TR = \sum_{i=1}^{S} t_i^0$ |
| Taxonomic Distinctness $\Delta^+$ | 1 (1) | Average path length between any two randomly chosen individuals (of different species) with the special case of only using presence/absence data for each species (Clarke and Warwick 1998). | $\Delta^+ = \left[\sum\sum_{i<j} \omega_{ij}\right] / [s(s-1)/2]$ |
| Hill's No. 2 | 1 (0) | Simpson's diversity converted into an 'effective number of species' that has the same units as species richness (Jost 2006). | $^2D = 1/\sum_{i=1}^{S} p_i^2$ |

Table continues on the next page.

[1] Number of studies that used the respective index.
[2] The bracketed number denotes the number of studies that exclusively used the respective index.

| | | | |
|---|---|---|---|
| $S_{PIE}$ | 1 (0) | Number of equally abundant species needed to yield the observed PIE (Chao et al. 2014). PIE is the probability that two individuals sampled randomly from a community are of different species (Hurlbert 1971). | $S_{PIE} = 1/(1 - PIE)$ <br> $PIE = 1 - \sum_{i=1}^{S} p_i^2$ |

**Note:** The information given derives from a total of 40 studies. Due to insufficient data for most indices, we continued this analysis with species richness and Shannon entropy only. In the final analysis, we only included 38 of the studies as 2 studies solely used indices that were not further evaluated.

**Abbreviations:** S = total number of species in a community; $p_i$ = relative abundance of species i, $t_i$ = relative abundance of taxon i; N = total number of individuals; $\omega_{ij}$ = 'distinctness weight' for the path length between species i and j.

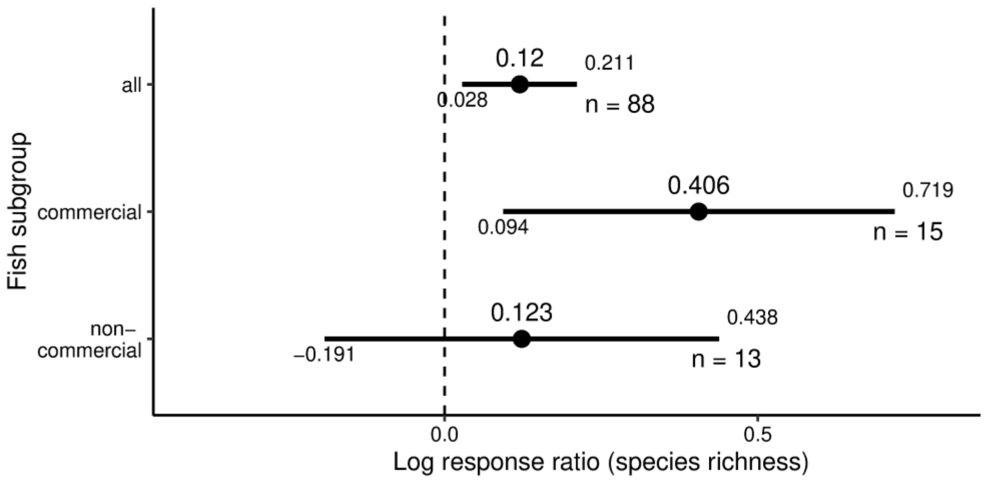


**Figure S1** Average log response ratios (MPA:unprotected) ± 95% confidence intervals as a function of fish group as all, commercially targeted fish, and non-commercial fish. The corresponding values can be found in Table 2: Model 2. A log response ratio of 0 indicates no effect through protection. Positive log response ratios indicate greater diversity inside the MPA boundaries relative to outside and negative values indicate greater diversity outside the MPA boundaries relative to inside. n values are the sample sizes of the mean estimates.

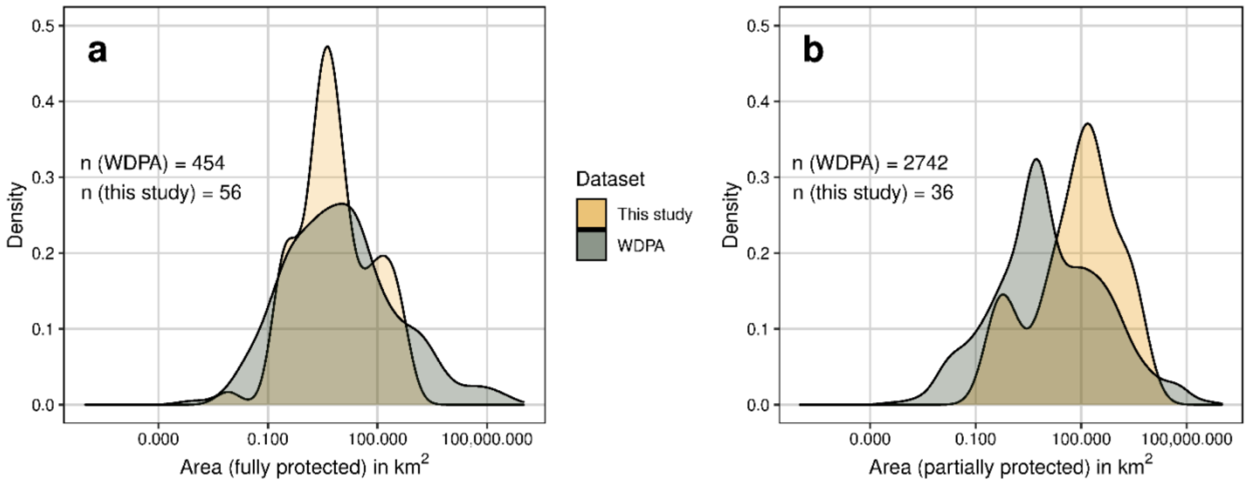


**Figure S2** Distribution of MPA size in $km^2$ in this study versus the World Database on Protected Areas (UNEP-WCMC and IUCN 2021) for (a) fully protected areas and (b) partially protected areas.

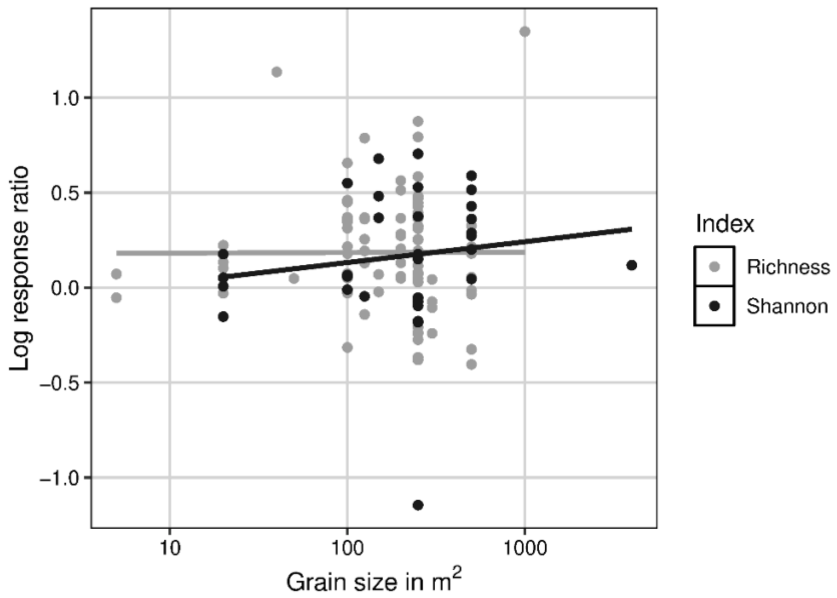


**Figure S3** Log response ratio (MPA:unprotected) of species richness and Shannon entropy as a function of grain of samples in $m^2$. There is no significant relationship between the grain of samples and the log response ratio of species richness (Spearman's rank correlation, S = 149495, $p > 0.05$) or the log response ratio of Shannon entropy (Spearman's rank correlation, S = 4212, $p > 0.05$).

### Meta-analytical Linear Mixed Models

To complement the linear mixed models, we fit meta-analytic linear mixed models using the 84% of studies that reported suitable error estimates. This analysis was weighted and gave studies with putative higher precision greater weight. For weighting we calculated the standard error of the log response ratio (Hedges, Gurevitch, and Curtis 1999) using:

$$se(lnRR) = \sqrt{\frac{{sd_{pa}}^2}{n_{pa}{\bar{x}_{pa}}^2} + \frac{{sd_{oa}}^2}{n_{oa}{\bar{x}_{oa}}^2}},$$

(Equation S1)

where $n_{pa}$ and $n_{oa}$ are the sample sizes of the mean estimates, $x_{pa}$ and $x_{oa}$ are the mean diversities at the protected site and unprotected site, respectively, and $sd_{pa}$ and $sd_{oa}$ are the associated standard deviations. Bayesian hierarchical models were fit using the Markov chain Monte Carlo (MCMC) sampling method and coded using the brms package (version 2.15.0, Bürkner 2017). All models were fit with four chains and 4,000 iterations, with 2,000 used as warmup. We used weakly informative priors (Williams et al. 2018) for μ and Half-Cauchy priors for τ (extraDistr package version 1.9.1, Wolodzko 2020). The only difference between the meta-analytic model and those presented in the main text was the incorporation of the standard error of the log response ratio, the meta-analytic models all took the same form to those described in Table 1. Convergence was verified using the potential scale reduction factor (PSRF) and visual inspection of the MCMC chains.

**Table S2** Comparison of the model results of the linear mixed models (lmm) versus the meta-analytic linear mixed models (meta-lmm).

| Model[3] | Response | Fixed Effect | Estimate | | Lower | | Upper | |
|---|---|---|---|---|---|---|---|---|
| | | | *lmm* | *meta-lmm* | *lmm.* | *meta-lmm* | *lmm* | *meta-lmm* |
| 1 | lnRR(SR) | Intercept | 0.169 | 0.15 | 0.097 | 0.06 | 0.243 | 0.24 |
| 2 | lnRR(SR) | MPA size | -0.046 | -0.02 | -0.073 | -0.02 | -0.017 | -0.02 |
| 3 | lnRR(SR) | MPA age | -0.018 | -0.06 | -0.104 | -0.06 | 0.063 | -0.05 |
| 4 | lnRR(SR) | Partial protection | -0.086 | -0.10 | -0.213 | -0.11 | 0.052 | -0.09 |
| 7 | lnRR(H) | Intercept | 0.126 | 0.13 | -0.016 | -0.05 | 0.266 | 0.33 |

**Note:** For the linear mixed models, 'Estimate' corresponds to mean log response ratios and for the meta-analytic linear mixed models 'Estimate' corresponds to median log response ratios. For the linear mixed models 'Lower' and 'Upper' correspond to 95% confidence intervals and for the meta-analytic linear mixed models to 95% credible intervals. Model results are presented in an abbreviated form, with only the fixed effects of interest shown.

**Abbreviations:** lnRR = log response ratio; SR = species richness; MPA size = mean-centered and log-transformed MPA size in $km^2$; MPA age = mean-centered and log-transformed MPA age in years; H = Shannon entropy

[3] Model numbers refer to the linear mixed models described in Table 1.

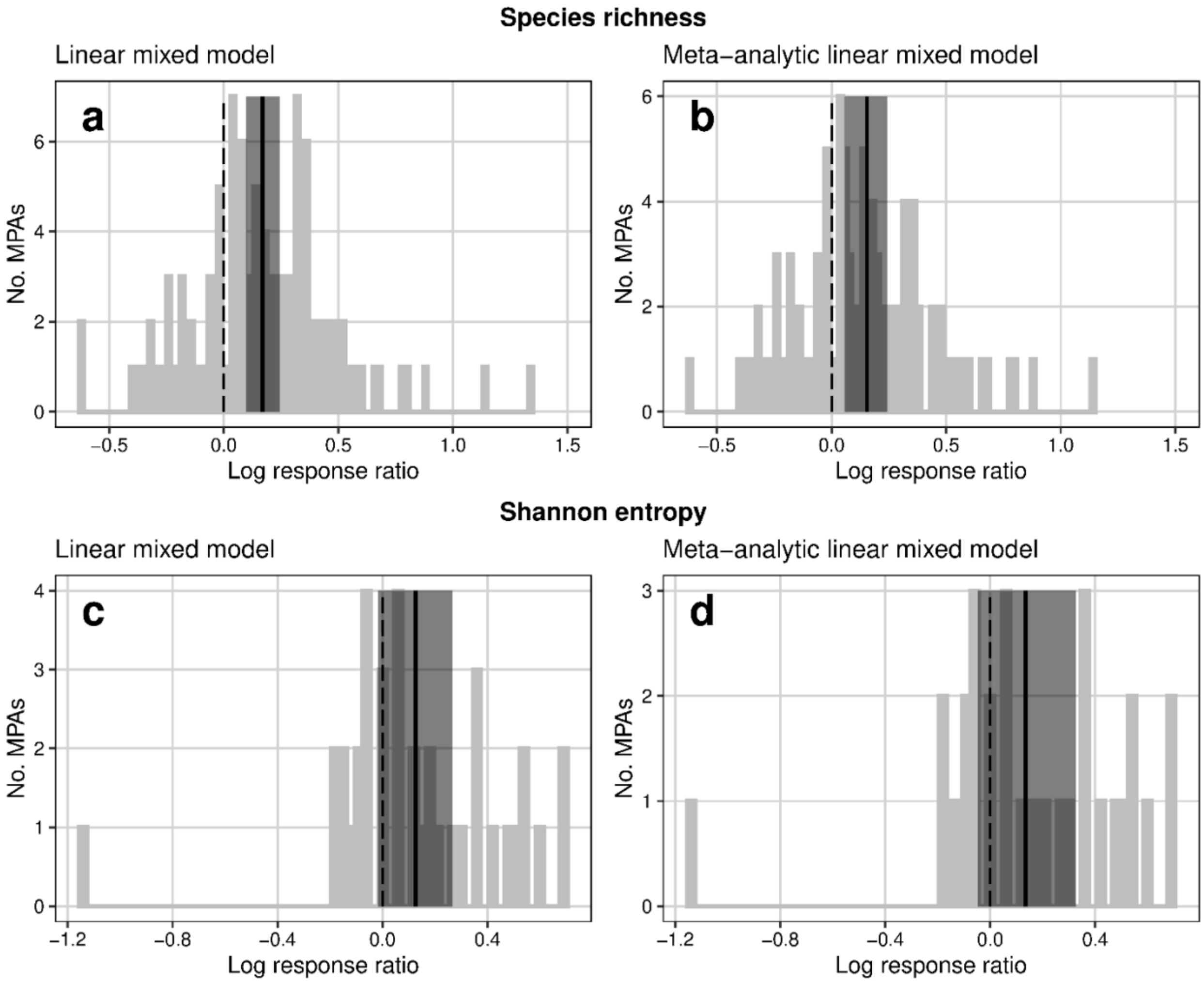


**Figure S4** Visual comparison of the results of the linear mixed models (a) and (c) versus the meta-analytic linear mixed models (b) and (d). All four subplots show the overall effect of protection on biodiversity as the distribution of log response ratios (MPA:unprotected) of ((a) and (b)) species richness and ((c) and (d)) Shannon entropy. In (a) and (c) solid black lines indicate the mean log response ratios and dark-grey shading 95% confidence intervals. In (b) and (d) solid black lines indicate the median log response ratios and dark-grey shading 95% credible intervals. The corresponding values can be found in Appendix S2: Table S2: Model 1 for (a) and (b) and Model 7 for (c) and (d). In all subfigures, a log response ratio of 0 indicates no effect through protection. Positive log response ratios indicate greater diversity inside the MPA boundaries relative to outside and negative values indicate greater diversity outside the MPA boundaries relative to inside. Sample sizes for (a), (b), (c), and (d) were 114, 92, 38 and 32, respectively.

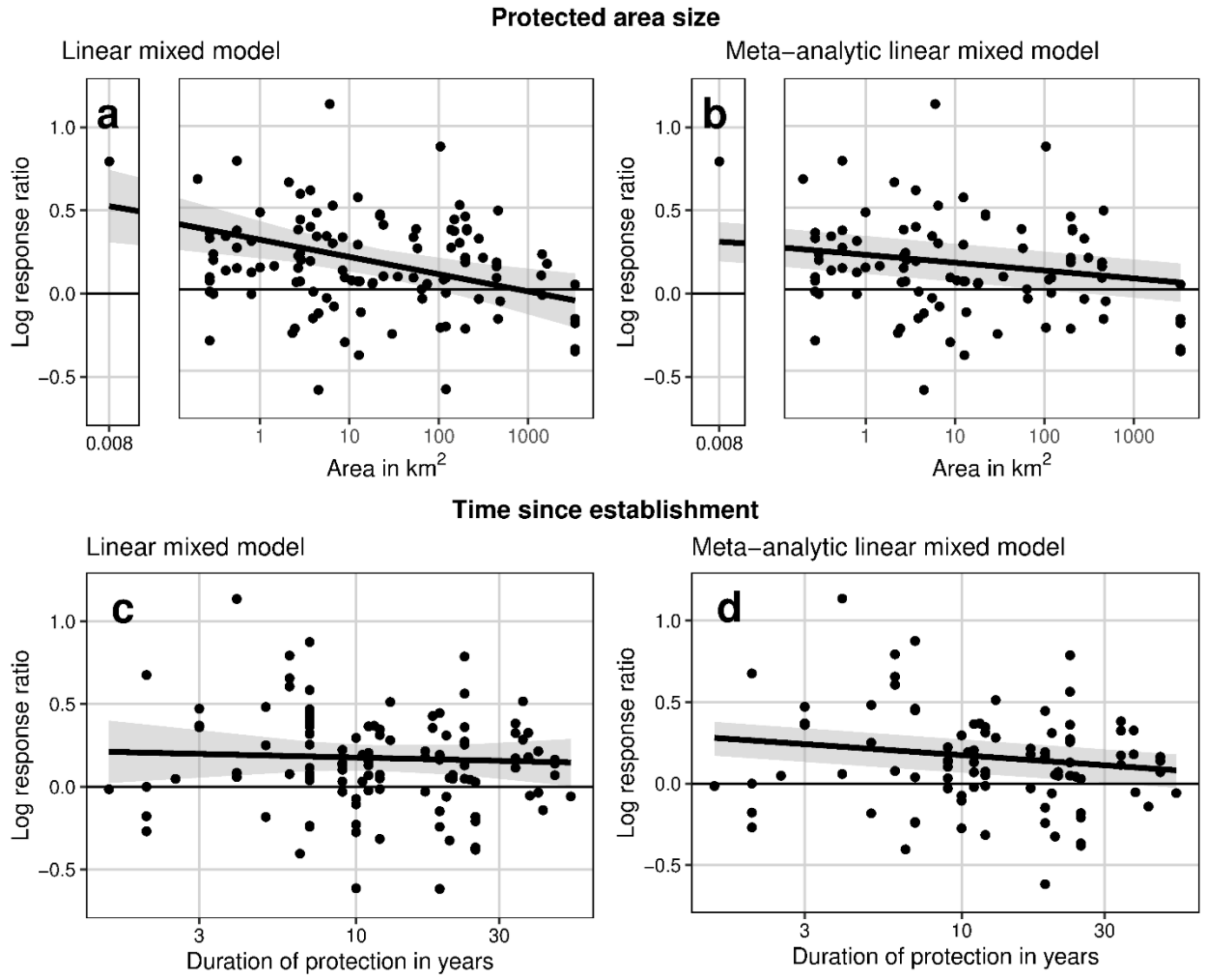


**Figure S5** Visual comparison of the results of the linear mixed models (a) and (c) versus the meta-analytic linear mixed models (b) and (d). Subplots (a) and (b) show the log response ratio (MPA:unprotected) of species richness as a function of MPA size in $km^2$ and (c) and (d) show the log response ratio (MPA:unprotected) of species richness as a function of MPA age as the number of years between MPA establishment and date of survey. In A and C grey shading indicates 95% confidence intervals and in B and D 95% credible intervals. The corresponding values can be found in Appendix S2: Table S2: Model 2 for (a) and (b) and Model 3 for (c) and (d). In both subfigures, a log response ratio of 0 indicates no effect through protection. Positive log response ratios indicate greater diversity inside the MPA boundaries relative to outside and negative values indicate greater diversity outside the MPA boundaries relative to inside. The sample size for (a) and (c) was 114 each and for (b) and (d) 92 each. Note: in (a) and (b), plots are split into two subplots each for better readability of the data.
